# Targeting the Untargetable: Predicting Pramlintide Resistance Using a Neural Network Based Cellular Automata

**DOI:** 10.1101/211383

**Authors:** Eunjung Kim, Ryan Schenck, Jeffrey West, William Cross, Valerie Harris, Joseph McKenna, Heyrim Cho, Elizabeth Coker, Steven Lee-Kramer Kenneth Y. Tsai, Elsa R. Flores, Chandler Gatenbee

## Abstract

De novo resistance is a major issue for the use of targeted anticancer drugs in the clinic. By integrating experimental data we have created a hybrid neural network/agent-based model to simulate the evolution and spread of resistance to the drug Pramlintide in cutaneous squamous cell carcinoma. Our model can eventually be used to predict patient responses to the drug and thus enable clinicians to make decisions regarding personalized, precision treatment regimes for patients.

## I Background

### Ia Cutaneous Squamous Cell Carcinoma

Cutaneous squamous cell carcinoma (CSCC) is a nonmelanoma skin cancer that, as of 2012, is estimated to affect 86,157 to 419,543 individuals in the United States, and has been described as a under-recognized health crisis [3]. The primary risk factor for CSCC is Ultraviolet (UV) light exposure, which can cause mutations that knock out p53 [1]. However, immunosuppression and infection with human papillomavirus, especially types 6, 11, 16, and 18, are also important risk factors for CSCC[1]. The prognosis for primary CSCC is very good, but the survival for metastatic CSCC is extremely low [1].

### Ib Targeting p53 with Pramlintide

p53, mutated in more than 50% of human cancers, is part of a larger family of genes known as the p53 family, including p63 and p73. Using genetic mouse models to understand how p63 function can be used to target p53 mutant tumors, we have identified Pramlintide, which is currently used to treat type 1 and type 2 diabetes, as such a compound. Indeed, we found that systemic delivery of this drug in p53-deficient mice resulted in rapid tumor regression [4]. This result is significant, as it allows p53 negative cells to be targeted, something that has previously not been achieved. However, resistance is an issue for the use of targeted anticancer drugs in the clinic. The goal of our project is to integrate experimental data with a computational model that will facilitate the identification of patients that will respond to Pramlintide.

## II Computational Model

### IIa Data

We have developed seven CSCC cell lines that are either sensitive and resistant to Pramlintide. The glycolytic activity of these cell lines, as measured using a Seahorse XF96 Bioanalyzer and a glycolysis stress assay, was significantly different between those resistant and sensitive to Pramlintide, where cells with high glycolytic activity were sensitive, while low glycolytic activity failed to exhibit a drug response. Xenograft models were treated with Pramlintide and their response correlated directly to glycolytic activity, again suggesting that glycolytic activity is related to sensitivity to Pramlintide.

**Table 1:**
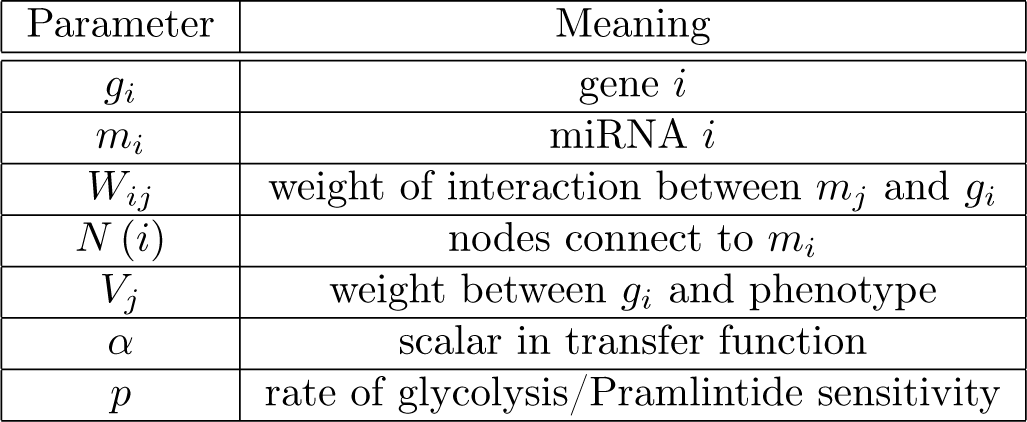
Parameters used in simulation

miRNA are involved in gene regulation, and so we next measured expression of miRNA associated with glycolysis (and thus Pramlintide efficacy) in each of the CSCC cell lines. Using the Wilcox Rank-Sum test (P-values <0.05), we we identified 30 differentially expressed microRNAs across the resistant and sensitive cell lines (Figure 1). Experimentally validated targets of the identified microRNAs was compiled using Diana Tools [5]. These candidate miRNAs were refined by cross-referencing their targets with proteins of the known glycolysis pathway (BioCarta Systematic Name M15109) [2]. Through this process we were able to identify miRNA that play an important role in altering the expression of genes involved in glycolysis, and thus Pramlintide sensitivity.

**Figure 1:**
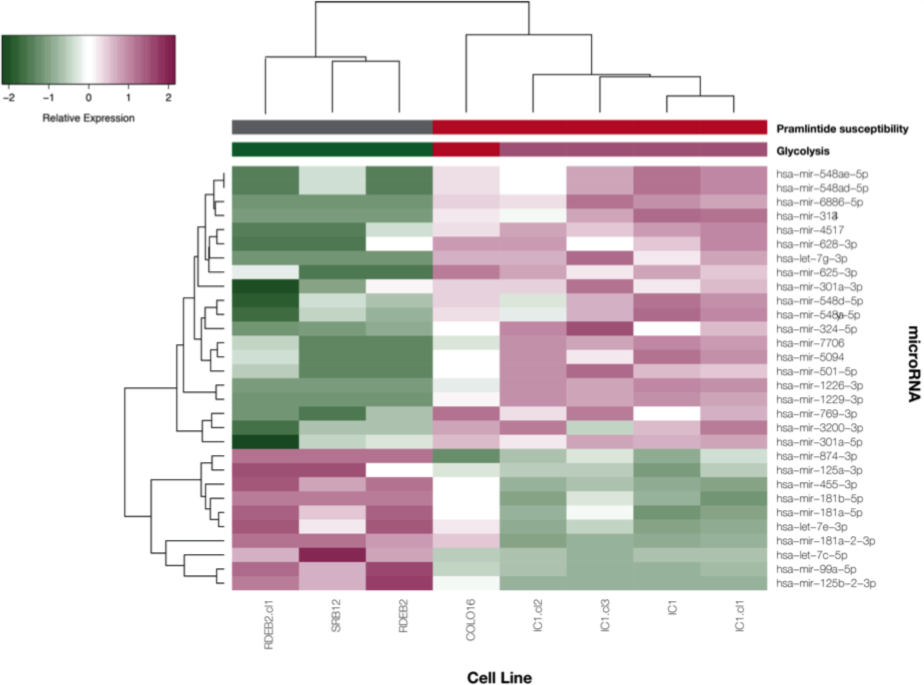
Expression profile of the 30 miRNA identified as being differentially expressed across susceptible and resistant cell lines

### IIb Neural Network

Understanding the relationship between miRNAs, genes, and phenotype is an overwhelming task considering the many levels of simultaneous interactions occurring in a growing cancer. Mathematical modeling may be a way forward with its ability to integrate multi-scale dynamics and to describe both linear and non-linear feedback between model agents. Here, we utilized a neural network modeling approach as a means to connecting miRNAs to genes and genes to phenotype

The above miRNA-gene-phenotype map was used to create a neural network that determines how the rate of glycolysis changes in response to miRNA expression (Figure 2). In this network, each of the 13 nodes in the first layer represent an miRNA, the middle layer are the genes identified to be targets of the miRNA, and the output layer is the phenotype, i.e. the rate of glycolysis/Pramlintide sensitivity. The links between the miRNA layer and gene layer were determined using the gene targets discovered using Diana Tools, as described above. Thus, the constructed neural network accurately represents the biological miRNA-gene-phenotype map.

**Figure 2:**
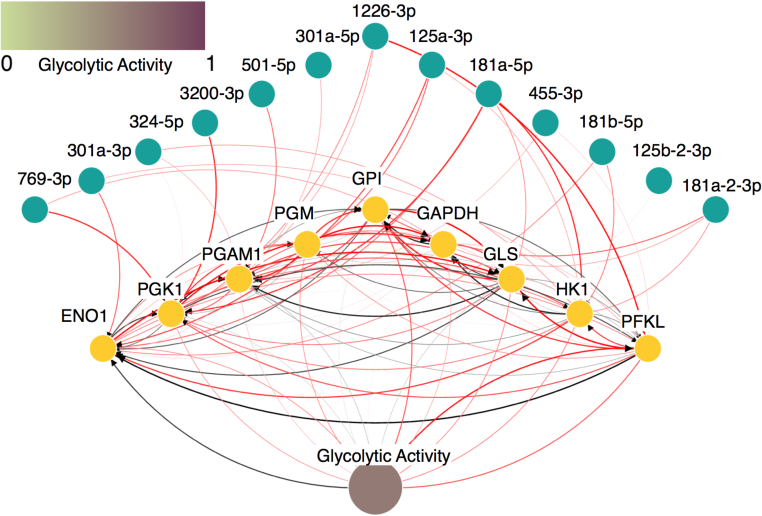
Neural network describing miRNA-genephenotype map

Given the structure of the neural network, each of the 13 miRNA, *m*_*i*_ =∈ [0, 1] = 1, 2,… 13, influence the activity of the associated transcripts, *g*_*i*_, in a linear additive way through a transfer function *g*_*i*_ = tanh (*α*∑_*j*∈*N*(*i*)_ *W*_*ij*_*m*_*j*_), where *W*_*ij*_ models the connections between miRNAs and genes, and *N* (*i*) is the set of nodes (either miRNAs or genes) connected to the miRNA *m*_*i*_. The phenotype, *p*, is modeled by *p* = tanh (*αV*_*j*_*g*_*j*_), where *V*_*j*_ is the connection strength between the genes and the phenotype. A Monte Carlo simulation is used to obtain weight matrices that recapitulate biological data.

### IIc Agent Based Model

The trained neural network is implanted into cells in a two dimensional on-lattice agent based model and is used to model phenotypes with varying rates of glycolysis and degrees of susceptibility to Pramlintide. As high glycolysis is associated with Pramlintide sensitivity, fitness is in inversely related to *p*, i.e. high sensitivity means lower fitness. After the cell uses the neural network to calculate *p*, it dies if *X ~ Binomial* (*p*) = 1. If the cell does not die, there is free adjacent space, and enough time has elapsed since the cell’s previous division, the cell will divide. Figure 3 for the decision tree used by cells in the model.

**Figure 3:**
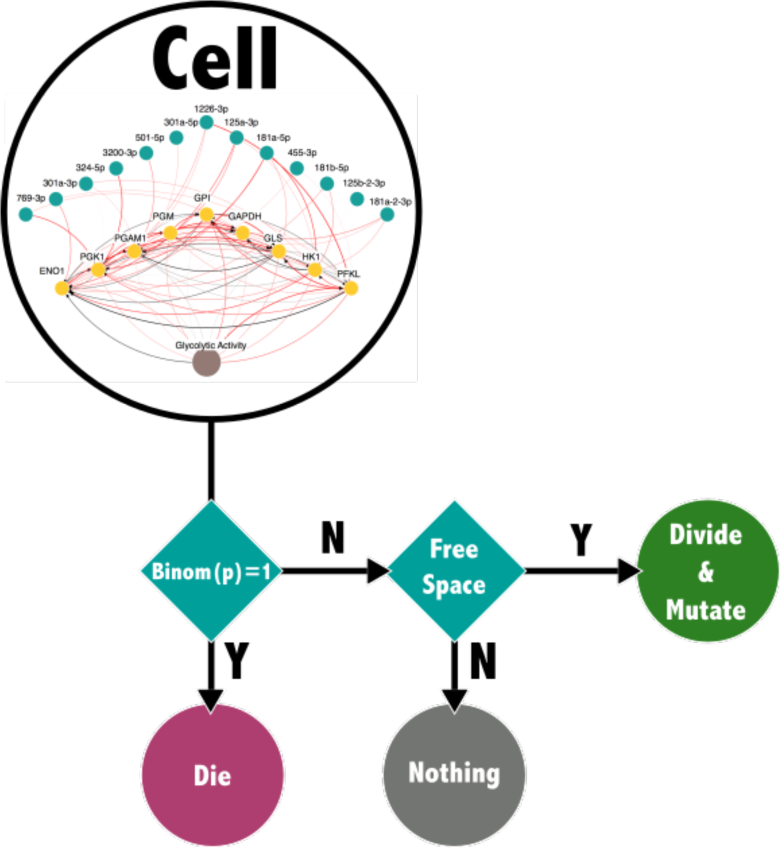
Decision tree for cells in agent based model

We simulate evolution by perturbing weights between miRNAs, genes, and glycolytic phenotype during cell division, thus allowing potential acquisition of resistance to Pramlintide. Similarly, the alteration of gene interactions will result in a new clone that will respond differently Pramlintide. Weights are perturbed using 2 methods: drift and mutations. The values of *W* and *V* are allowed to drift from the base network (trained via the neural network methods described above) by a difference of a Gaussian distribution random variable (mean = 0, *σ* = 0.001). Mutations occur during division with small probability (μ = 0.0000524), which selects a random column of W to mutate (i.e. set that column to Gaussian; mean = 0, *σ* = 1). Over time, this process will generate a heterogeneous population of clones with varying degrees of resistance to Pramlintide.

## III Application of Methods

We use our embedded neural network agent-based model to simulate the expansion of a single clone in the presence of Pramlintide. As the the tumor grows, there will be selection for clones that evolve resistance to treatment. Given that the weights embedded into the initial clone are user defined, it is possible initialize the simulation using a cell that has a patient’s miRNA expression profile. Conducting the patient specific simulation multiple times will generate a distribution of times until the evolution of drug resistance. This distribution can then be used to by the clinician as an additional tool to help them decide whether or not the patient is a good candidate from Pramlintide.

Figure 4 shows three sample simulations: a simulation administering Pramlintide without initial resistance (row 1), without initial resistance but accruing resistance de novo (row 2) and with a small fraction (1%) of resistant cell lines (row 3). The first case shows good results for Pramlintide treatment (the tumor is completely killed). The second shows a relapse, selecting for low glycolytic resistant cell lines that have accrued via random mutations and drift (blue cells). The third row shows a faster relapse due to the initial presence of resistant cell lines.

**Figure 4:**
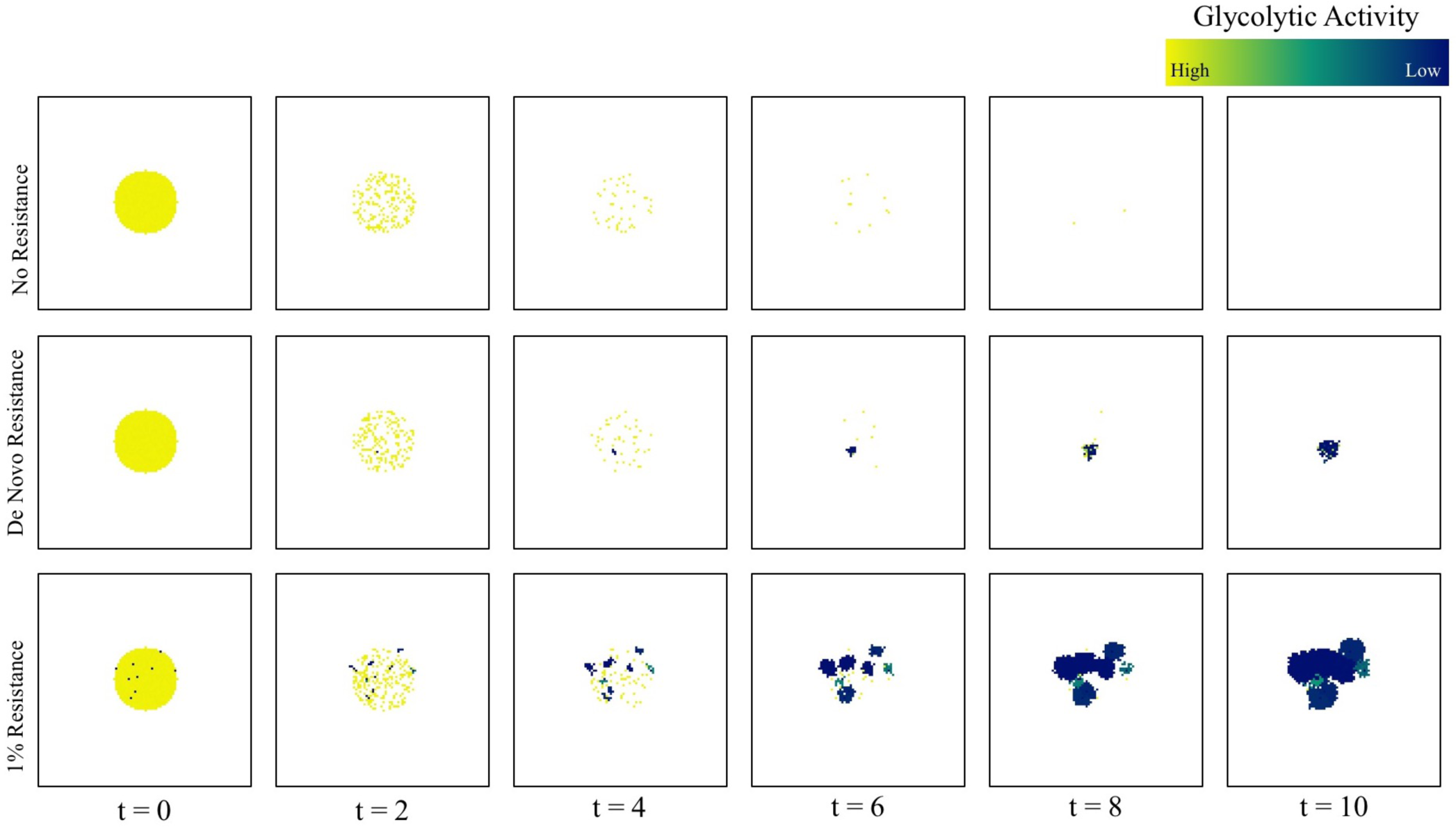
Sample of simulation runs.

## IV Closing Remarks

By leveraging a variety of tools, we were able to identify 13 miRNA, and their target genes in the glycolytic pathway, to construct a biologically defined neural network than can be embedded into cells within an agent based model. By running this simulation multiple times using a patient’s miRNA profile in the initial cell, a clinician can get a sense of how well a patient may respond to treatment. It is our hope that this model can thus serve as an additional tool for the clinician to use in making treatment decisions.

## Acknowledgements

We would like to thank the IMO Chair, Dr. Alexander Anderson, for organizing the 6th Annual Moffitt IMO workshop: Resistance, where this project was conceived. We are also extremely grateful to the Moffitt Cancer Center and the Moffitt PSOC for supporting this workshop through the NCI U54CA193489 grant.

